# Pheno4J: a gene to phenotype graph database

**DOI:** 10.1101/142257

**Authors:** Sajid Mughal, Ismail Moghul, Jing Yu, Tristan Clark, David S Gregory, Nikolas Pontikos

## Abstract

**Summary:** Efficient storage and querying of large amounts of genetic and phenotypic data is crucial to contemporary clinical genetic research. This introduces computational challenges for classical relational databases, due to the sparsity and sheer volume of the data. Our Java based solution loads annotated genetic variants and well phenotyped patients into a graph database to allow fast efficient storage and querying of large volumes of structured genetic and phenotypic data. This abstracts technical problems away and lets researchers focus on the science rather than the implementation. We have also developed an accompanying webserver with end-points to facilitate querying of the database.

**Availability and Implementation:** The Java code and python code is available at https://github.com/phenopolis/pheno4i

## 1 Introduction

A recurring theme in clinical genetics is to annotate large numbers of genetic using the Variant Effect Predictor (VEP) (McLaren et al. 2016) variants from well phenotyped patients, and load them into a database for efficient querying and filtering. However, a sequenced human genome typically produces more than 4 million genetic variants per individual, at least 100,000 of which are novel. This introduces a number of challenges for the conventional relational table based database model, which are better handled by graph databases.

The first challenge for a relational database is the efficient storage and querying of many-to-many relationships such as genetic variant to individual relationships. In a relational database, genetic variants and individuals would typically be stored as rows in two distinct tables and, in order to link them, a third table, which would be a very large mapping table known as a ‘join’ table, would need to be queried. While workable with small relational databases, ‘join’ queries quickly become inefficient as the number of relationships increases. On the other hand, in a graph database, data is stored in a manner such that ‘join’ queries are not required. Instead of using tables, each data record is stored as a distinct node with added relationships linking nodes stored internally as pointers. As such, analysing the relationship between nodes representing individuals and nodes representing genetic variants, is as simple as finding the connected nodes. This implies that query time remains constant despite a growing number of relationships. Additionally, by supporting multiple types of relationships that can be labelled, graph databases have an intuitive schema (Figure 1).

**Figure 1:**
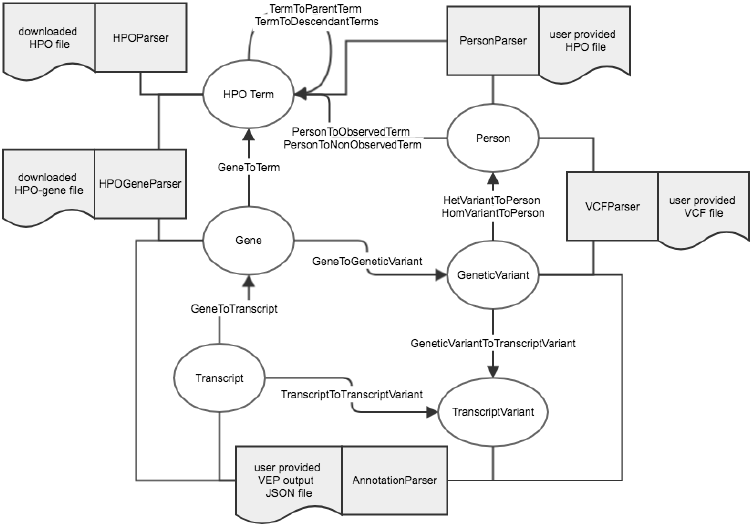
Overview of the Pheno4J graph database design. The grey boxes represent the five input files and five file parsers that are used to produce the graph database. The ellipses represent the six nodes types defined and the arrows represent the relationship types between the nodes.

The second challenge for a relational database is the extensibility of the database schema. Since each genetic variant is associated with an increasing number of annotation sources, which tend to be sparse and not always consistently formatted, the schema of relational database would have to be redefined every time a new annotation source is added. In a graph database, the schema is dynamically extensible to accommodate new sources of information by adding new types of nodes, node attributes or relationships (see Supplementary Section 3).

Finally, directed acyclic graph ontologies such as the Human Phenotype Ontology (HPO) (Robinson et al. 2008) and the Gene Ontology can be directly stored and queried in graph databases whereas complex operations would be required to achieve the same in a relational database.

In order to address these challenges, we have developed Pheno4J (https://github.com/phenopolis/pheno4j/), a tool implemented in Java that parses, integrates and imports genotype, annotated genetic variants and patient phenotype files into a Neo4J graph database. Using the Cypher querying language, it is then possible to perform sophisticated queries in real-time (Supplementary Section 2). In our live installation, we have loaded 5,025 exomes and 4M variants. This amounts to 8M nodes, 487M relationships and 296M properties; which when stored in memory takes up approximately 20 Gb of memory (runtime 40 minutes). The scalability of our solution with respect to the number of exomes stored has been demonstrated in Supplementary Section 4.

## 2 Implementation

In order to build the database, a total of five data files are required as input. These include three user generated files and two publically available downloadable files (Figure 1). Trimmed down versions of these files have been included in the GitHub repository for testing purposes. The three user generated input files are:

- VCF file containing the person to genetic variant relationships.
- JSON generated by the VEP containing the annotation for each variant. This produces the genetic variant-to-gene, transcript-to-gene and genetic variant-to-transcript relationships.
- Phenotype CSV file containing the link from persons to HPO terms.

The other two input file are publicly downloadable:

- The HPO ontology which is obtained automatically from the HPO website.
- The gene to HPO file that can be downloaded from the HPO website.

These files are parsed and then loaded into the database. Supplementary Section 1 shows the steps required for building and running the database.

## 3 Use cases

Once the database is loaded, the data can then be queried using the Cypher language. One basic application could be to identify rare damaging variants by filtering by frequency and Combined Annotation Dependent Depletion (CADD) score (Kircher et al. 2014). For example, returning all variants that have a frequency less than 0.001 and a CADD score greater than 20, yields 171,532 variants from our cohort of 6,467 exomes (runtime 2.6 seconds). In Cypher this would be:

~~~
MATCH (gv:GeneticVariant)
WHERE gv.cadd_phred > 20 AND
gv.allele_freq < 0.001 AND gv.kaviar_AF < 0.0001
RETURN count(gv.variantId);
~~~

Another application could be to identify related individuals by counting the number of rare heterozygous variants shared with “person1”. Here we return the list of ten individuals by decreasing shared rare variant count (runtime 1.2 seconds).

~~~
MATCH (k:Person)
WITH count(k) as numberOfPeople
MATCH (p:Person {personId:”person1”})<-[:HetVariantToPerson]-
(gv:GeneticVariant)
WHERE gv.allele_freq < 0.001 AND gv.kaviar_AF < 0.001
WITH size(()<-[:HetVariantToPerson]-(gv)) as het_count, gv, p,
numberOfPeople
WHERE het_count > 1 AND ((het_count/toFloat(numberOfPeople)) <= 0.05)
 // Sharing of variants with Person q.
MATCH (gv)-[:HetVariantToPerson]->(q:Person)
WHERE p <> q
WITH p,q,count(gv) as c
ORDER BY c desc LIMIT 10
RETURN p.personId,q.personId, c;
~~~

In this query, we make use of Cypher’s efficient edge counting `size(()<-[:HetVariantToPerson]-(gv)) as het_count` to get the number of outgoing relationship of a node.

Another use case might be to find all individuals with a given HPO term such as for example “Retinal dystrophy” (HP:0000556), which returns 521 individuals in our cohort (runtime 0.3 seconds). This query returns all individuals including those that might be annotated with a child term (direct or indirect) of “Retinal dystrophy” such as “macular dystrophy”.

~~~
MATCH (p:Term {name:’Retinal dystrophy’})-
[:TermToDescendantTerms]->(q:Term)<-[:PersonToObservedTerm]-
(r:Person)
RETURN r.personId;
~~~

In order to obtain a list of candidate variants, we can query all rare damaging homozygote variants seen in people with “Retinal dystrophy” that belong to a known “Retinal dystrophy” gene. In our data, this returns 69 distinct variant ids (runtime <1 second).

~~~
MATCH (p:Term {name:’Retinal dystrophy’})-
[:TermToDescendantTerms]->(q:Term)<-[:GeneToTerm]-(gs:Gene)
WITH distinct gs as retinal_dystrophy_genes
~~~

~~~
MATCH (retinal_dystrophy_genes)-[:GeneToGeneticVariant]-
>(gv:GeneticVariant)
WHERE gv.allele_freq < 0.001 AND gv.cadd_phred > 20 AND
gv.kaviar_AF < 0.001
WITH distinct gv, retinal_dystrophy_genes
~~~

~~~
MATCH (r:Person)<-[:HomVariantToPerson]-(gv)
WITH distinct gv, retinal_dystrophy_genes, r
~~~

~~~
MATCH (p:Term {name:’Retinal dystrophy’})-
[:TermToDescendantTerms]->(q:Term)<-[:PersonToObservedTerm]-(r)
~~~

~~~
RETURN distinct r.personId, gv.variantId,
retinal_dystrophy_genes.gene_name;
~~~

Conversely, we can suggest “Retinal dystrophy” as a phenotype for individuals that have rare damaging homozygote variants in recessive retinal dystrophy genes:

~~~
MATCH (p:Term)-[:TermToDescendantTerms]->(q:Term)<-
[:PersonToObservedTerm]-(r:Person)
WHERE p.name=’Retinal dystrophy’
WITH COLLECT(distinct r) as persons
MATCH (p:Person) WHERE NOT p IN persons WITH COLLECT(p) as
non_retinal_dystrophy_persons_list
~~~

~~~
MATCH (t:Term)<--(g:Gene)-->(t2:Term) WHERE t.name=’Retinal
dystrophy’ AND t2.name=’Autosomal recessive inheritance’
WITH COLLECT(DISTINCT g) as
recessive_retinal_dystrophy_genes_list,
non_retinal_dystrophy_persons_list
~~~

~~~
UNWIND recessive_retinal_dystrophy_genes_list as
recessive_retinal_dystrophy_genes
MATCH (recessive_retinal_dystrophy_genes)-->(gv:GeneticVariant)
WHERE gv.allele_freq < 0.001 AND gv.cadd_phred > 25 AND
gv.kaviar_AF < 0.0001 WITH gv,
non_retinal_dystrophy_persons_list
~~~

~~~
UNWIND non_retinal_dystrophy_persons_list as
non_retinal_dystrophy_persons
MATCH (recessive_retinal_dystrophy_genes)-->(gv)-
[:HomVariantToPerson]->(non_retinal_dystrophy_persons)
RETURN distinct gv.variantId, gv.most_severe_consequence,
gv.cadd_phred, gv.kaviar_AF,
recessive_retinal_dystrophy_genes.gene_name,
non_retinal_dystrophy_persons.personId ORDER BY gv.cadd_phred
DESC;
~~~

Further queries and analysis are described in the Supplementary Section 2.

## 4 Discussion

Currently, there are bespoke tools such as GQT (Layer et al. 2016) and BGT (Li 2016), which are very space efficient and excel at querying large annotated VCFs in real-time.

However, they do not build on existing database technologies making them harder to extend and do not support HPO querying. At the other end of the spectrum, there are also solutions such as hail.is (https://github.com/hail-is/hail) for very large genomic datasets that are distributed over a cluster of computers.

However, these require access to significant infrastructure and the overheads of installation are not trivial. Our solution requires minimal programming and is particularly suitable to a web front end for interrogating the data of an exome database of around 10,000 well phenotyped individuals as typical in rare disease groups such as BRIDGE (https://bridgestudy.medschl.cam.ac.uk) and the UK Inherited Retinal Disease Consortium. We are using Pheno4J as a backend for our Phenopolis (https://phenopolis.github.io/) platform to replace our current NoSQL MongoDB. While still relatively in their infancy, we predict graph databases will become pervasive in biology with a growing number of projects adopting them (Pep Tracker DB (https://www.peptracker.com), OwlSim (www.owlsim.org) and SciGraph (https://github.com/SciGraph/SciGraph)).

## Acknowledgements

We also acknowledge the Computer Science High Performance Cluster for providing us with the computing platform on which to analyse our data. SM wrote the code. SM, JY, IM and NP wrote the manuscript. TC and DSG provided the computing infrastructure and the server configuration.

## References

Kircher, Martin, Daniela M. Witten, Preti Jain, Brian J. O’Roak, Gregory M. Cooper, and Jay Shendure. 2014. “A General Framework for Estimating the Relative Pathogenicity of Human Genetic Variants.” Nature Genetics 46 (3): 310–15.

Layer, Ryan M., Neil Kindlon, Konrad J. Karczewski, Exome Aggregation Consortium, and Aaron R. Quinlan. 2016. “Efficient Genotype Compression and Analysis of Large Genetic-Variation Data Sets.” Nature Methods 13 (1): 63–65.

Li, Heng. 2016. “BGT: Efficient and Flexible Genotype Query across Many Samples.” Bioinformatics 32 (4): 590–92.

McLaren, William, Laurent Gil, Sarah E. Hunt, Harpreet Singh Riat, Graham R. S. Ritchie, Anja Thormann, Paul Flicek, and Fiona Cunningham. 2016. “The Ensembl Variant Effect Predictor.” Genome Biology 17 (1): 122.

Robinson, Peter N., Sebastian Köhler, Sebastian Bauer, Dominik Seelow, Denise Horn, and Stefan Mundlos. 2008. “The Human Phenotype Ontology: A Tool for Annotating and Analyzing Human Hereditary Disease.” American Journal of Human Genetics 83 (5): 610–15.

